# A-to-I mRNA editing recodes CqsA and affects T6SS-mediated killing in *Vibrio*

**DOI:** 10.64898/2026.06.08.730849

**Authors:** Rinat Cohen-Pavon, Chaya Mushka Fridman, Danielle Arad, Sahar Melamed, Irina Rostovsky, Neta Sal-Man, Liam Aspit, Dor Salomon, Dan Bar Yaacov

## Abstract

Adenosine-to-inosine (A-to-I) RNA editing alters genetic information post-transcriptionally, yet its ecological and evolutionary significance in bacteria remains largely unexplored.

Here, we show that endogenous RNA editing has functional consequences in bacteria. Using *Vibrio alginolyticus* as a model, we identified 38 editing events—the highest number reported for any bacterium to date. Editing frequencies varied across growth phases and occurred within a shorter conserved sequence motif than observed in other bacteria, suggesting species-specific determinants. The mRNA of the quorum-sensing (QS) autoinducer synthase *cqsA* was the most extensively edited, with 70–90% of transcripts modified. Phylogenetic and experimental analyses revealed that *cqsA* editing is evolutionarily conserved across diverse *Vibrio* species, including human pathogens. Protein mass spectrometry showed that editing replaces a tyrosine with a cysteine residue at position 193 of endogenously expressed CqsA without altering its expression or the canonical downstream QS signaling pathway. However, we found that endogenous editing of *cqsA* alters the expression of a subset of genes and is required for efficient type VI secretion system (T6SS)–mediated interbacterial killing. Together, these findings suggest that CqsA has additional roles beyond its canonical QS function and that RNA editing can modulate bacterial physiology.

**Significance:** Bacteria are haploid organisms having a single copy of each gene. A-to-I RNA editing can change genetic information at the RNA level, creating protein isoforms in bacteria, but its functional impact remains unclear. Here we show that a marine *Vibrio* species has at least 38 edited RNAs—the highest number reported in any bacterium. We further show that editing of the quorum-sensing synthase *cqsA* is widespread across *Vibrio* species and recodes CqsA protein sequence without affecting canonical quorum sensing. Instead, editing alters expression of a focused gene set, including the type VI secretion system component *hcp1*, and is required for efficient T6SS-mediated interbacterial killing. Our findings uncover a conserved, quorum-sensing–independent role for CqsA in bacterial competition.

## Introduction

Adenosine-to-inosine (A-to-I) RNA editing is a post-transcriptional modification that changes the genetic information at the RNA level (1). In this process, adenosine is enzymatically deaminated into inosine, which is read as guanosine by the ribosome or genetic machineries (*e*.*g*., reverse transcriptase). Subsequently, A-to-I RNA editing can recode protein sequence and function (2, 3). In eukaryotes, A-to-I mRNA editing is catalyzed by adenosine deaminases acting on RNA (ADARs) and plays critical roles in neuronal function, immune regulation, and proteome diversification (1, 2, 4-17). Although initially thought to be exclusive to multicellular organisms, A-to-I mRNA editing has previously been identified in bacteria (18).

In bacteria, A-to-I RNA editing is mediated by the enzyme tRNA-specific adenosine deaminase (TadA), which was originally believed to target position 34 at tRNA-Arg2 exclusively (19). However, recent studies have demonstrated that TadA can also edit mRNA, leading to sequence modifications that may influence bacterial physiology (18, 20). While the full extent and functional consequences of A-to-I mRNA editing in bacteria remain poorly understood, emerging evidence suggests that it may contribute to key cellular processes, including stress responses, regulating toxin activity, motility, and virulence (20-22). Recently, we showed that A-to-I mRNA editing occurs in dozens of gammaproteobacterial species (23-25). However, our analysis included only publicly available RNA-sequencing (RNA-seq) data. We applied a series of stringent computational filters to reduce technical noise (*i*.*e*., false-discovery sites detection). However, we hypothesize that due to the stringent nature of our analysis, we underestimated the number of A-to-I editing events in the examined species. Thus, an in-depth analysis of corresponding DNA and RNA samples from various species is needed. Subsequently, identifying and characterizing A-to-I RNA editing events in bacterial species could provide insights into novel regulatory mechanisms that shape bacterial adaptation and pathogenesis.

*Vibrio* species are a diverse group of marine bacteria, many of which are known or emerging pathogens of humans and marine organisms (26-28). *Vibrio* species inhabit a wide range of aquatic niches, from coastal waters to estuarine and open-ocean environments, where they encounter fluctuating conditions. Despite extensive research on *Vibrio* species, the post-transcriptional mechanisms that help these bacteria adapt and respond to their environment remain poorly understood. In particular, RNA-based processes can rapidly modulate protein function without altering the underlying genome. For example, small RNAs were shown to modulate motility, quorum sensing (QS), and biofilm formation in *Vibrio cholerae* (29, 30). Moreover, RNA modifications were shown to be important for *Vibrio* biology (31). Exploring these RNA regulatory layers can therefore reveal additional strategies by which *Vibrio* species adapt, persist, and cause disease. Thus, an in-depth analysis of RNA editing, its occurrence, regulation, and function in *Vibrio* species is warranted.

*V. alginolyticus* has emerged as a major pathogen of aquaculture species and a frequent cause of human vibriosis, making it highly relevant for studying virulence regulation, quorum sensing, secretion systems, and host–pathogen interactions (32-34). Moreover, we identified endogenous A-to-I mRNA editing in *V. alginolyticus*, establishing it as an excellent model for investigating A-to-I editing in *Vibrio* species (23).

In this study, we investigate the occurrence and functional significance of A-to-I RNA editing in *Vibrio alginolyticus*. Using high-throughput sequencing, we identify novel mRNA editing events, examine how they change during bacterial growth, and define their regulatory determinants. We focus on an editing event in *cqsA*, showing that it is evolutionarily conserved across *Vibrio* species and important for T6SS activity. Together, our findings provide the first evidence that endogenous A-to-I RNA editing can shape bacterial physiology and behavior.

## Results

### The A-to-I RNA editing landscape of *V. alginolyticus*

First, we aimed to determine the A-to-I mRNA editing landscape across growth phases. To this end, we sequenced RNA and DNA from *V. alginolyticus* at the early-log, mid-log, and stationary phases to identify A-to-G RNA-DNA mismatches that could represent A-to-I editing events (Figure 1A-C). Examining the sequence around the A-to-G mismatches revealed that they are significantly enriched in a UACG sequence motif required by TadA for its activity (Figure 1D, Supplementary Figure 1, and Supplementary Table 1). Most UACG sites occur in coding sequences (CDS), and some occur in RNA mapped to intergenic regions or the antisense strand of known genes (Figure 1E and Supplementary Table 3). We also detected the canonical editing event at A34 in tRNA-Arg2 transcribed from six different genes with identical sequences (Figure 1E and Supplementary Tables 1-3). We further observed that editing events within CDS are predicted to alter protein sequence in most cases (Figure 1F and Supplementary Table 3). In total, we identified 38 UACG embedded A-to-I RNA editing sites in addition to the canonical tRNA-Arg2 edited site (Figure 1G and Supplementary Tables 2-3). Among the 38 sites, most (29) are novel and were not previously reported. Editing levels vary between transcripts and range between an average of 0.4% in *infB* to 82.62% in *cqsA*, across all RNA samples (Figure 1G and Supplementary Table 2). Notably, in some transcripts, editing was detected in only a subset of analyzed samples (*e*.*g*., *infB*), whereas in others it was detected in all samples (*e*.*g*., *cqsA*). To validate our identified editing events, we used Sanger sequencing of corresponding DNA and RNA samples (extracted from the same sample) of six sites with average editing levels above 15% (Figure 1G and Supplementary Table 2). Indeed, we observed the edited allele at all of the examined sites, validating our high-throughput RNA-seq computational analysis (Figure 1H).

**Figure 1.**
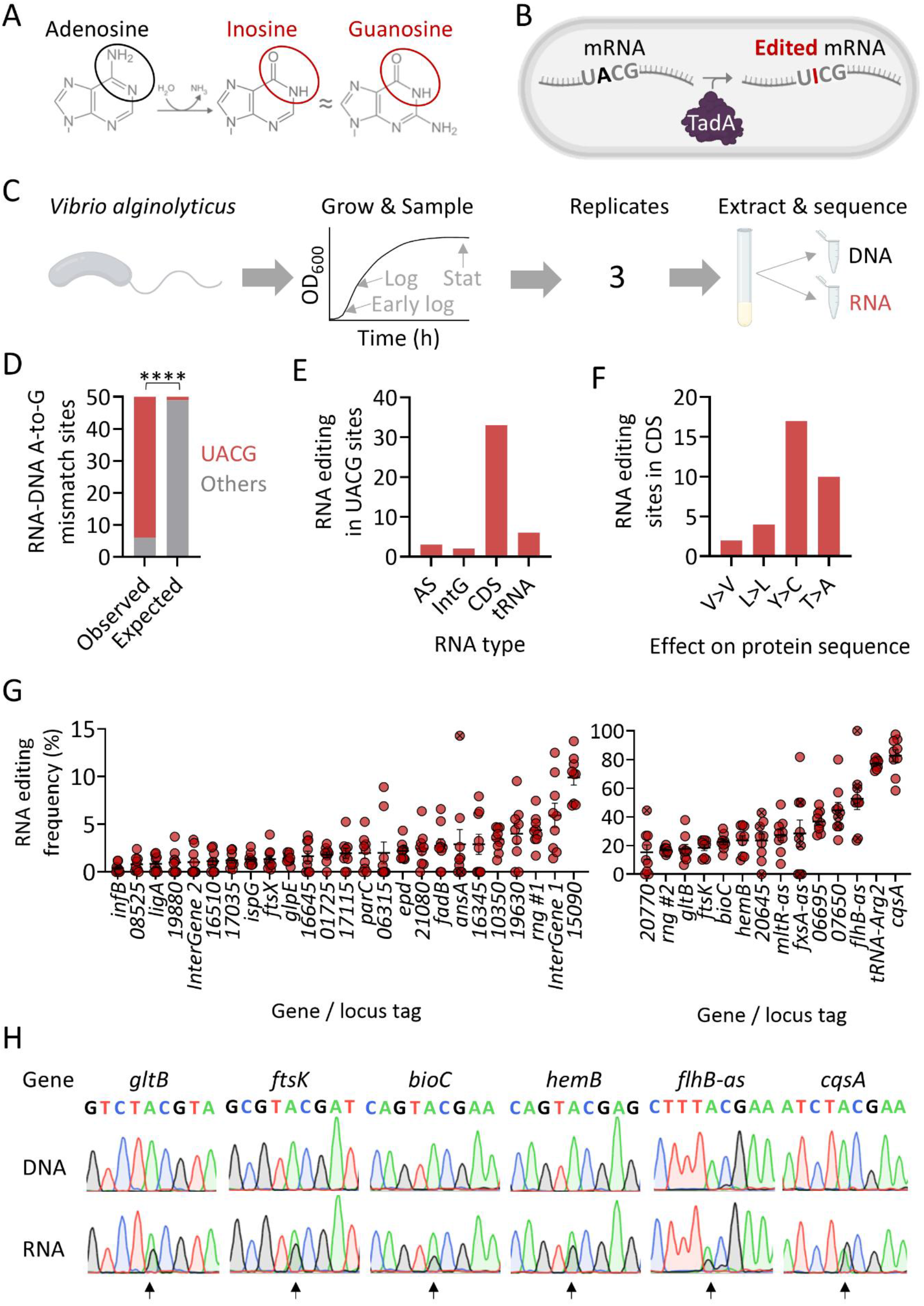
The landscape and levels of A-to-I RNA editing in *V. alginolyticus*. **A**. Adenosine is deaminated into inosine, which is recognized as guanosine by the ribosome and reverse transcriptase, allowing its detection in sequencing data. **B**. TadA can edit mRNAs containing a four-base sequence motif of UACG (TACG at the DNA level). **C**. Experimental design: *V. alginolyticus* was grown at 37 °C in Marine Broth (MB). RNA and DNA were extracted from the same sample at the early logarithmic, mid-logarithmic, and stationary phases, and RNA-seq and DNA-seq were performed. The experiment was performed with three biological replicates (we repeated the entire experiment three times on different days). **D**. RNA-DNA A-to-G mismatches are significantly enriched in UACG motif (Fisher’s exact test; p < 0.0001). **E**. Distribution of editing events according to RNA type. AS – antisense RNA; IntG – intergenic; CDS – coding sequence; Trna – tRNA-Arg2. **F**. Predicted effect of RNA editing on protein sequence (when in CDS). **G**. An overview of RNA editing levels in identified transcripts (mean and standard errors). On the left panel are sites with an average editing of below 15%. On the right panel are sites with an average editing of above 15%. Samples marked with “x” have read coverage lower than 10 reads. **H**. Sanger sequencing validates the occurrence of six editing events in corresponding DNA and RNA samples. The edited site is marked with a black arrow, and the editing signal is represented by a guanosine (black) peak in the RNA samples. The genomic positions of the edited sites are found in Supplementary Tables 1 and 2.

We conclude that our analysis expanded the repertoire of *bona fide* A-to-I RNA editing sites in *V. alginolyticus*, making it the bacterium with the most editing events reported to date.

### A-to-I mRNA editing in *V. alginolyticus* changes across growth phases in specific transcripts

Next, we sought to determine whether RNA editing levels in *V. alginolyticus* vary during different growth phases. To this end, we analyzed the identified RNA editing events with respect to their corresponding growth phase (Figure 1C and 1G). We observed that the average editing level of all genes, when examined as a group, was not significantly different between growth phases (Supplementary Figure 2). However, we hypothesized that significant changes in editing levels could be observed in a subset of transcripts, as we showed recently in *E. coli (24)*. In line with this hypothesis, we identified specific genes whose editing levels differed markedly between growth phases (Figure 2 and Supplementary Table 4). These findings suggest that, although editing levels across the transcriptome are generally only weakly coordinated, some genes exhibit phase-specific regulation of RNA editing, indicative of a gene-specific mechanism controlling editing levels.

**Figure 2.**
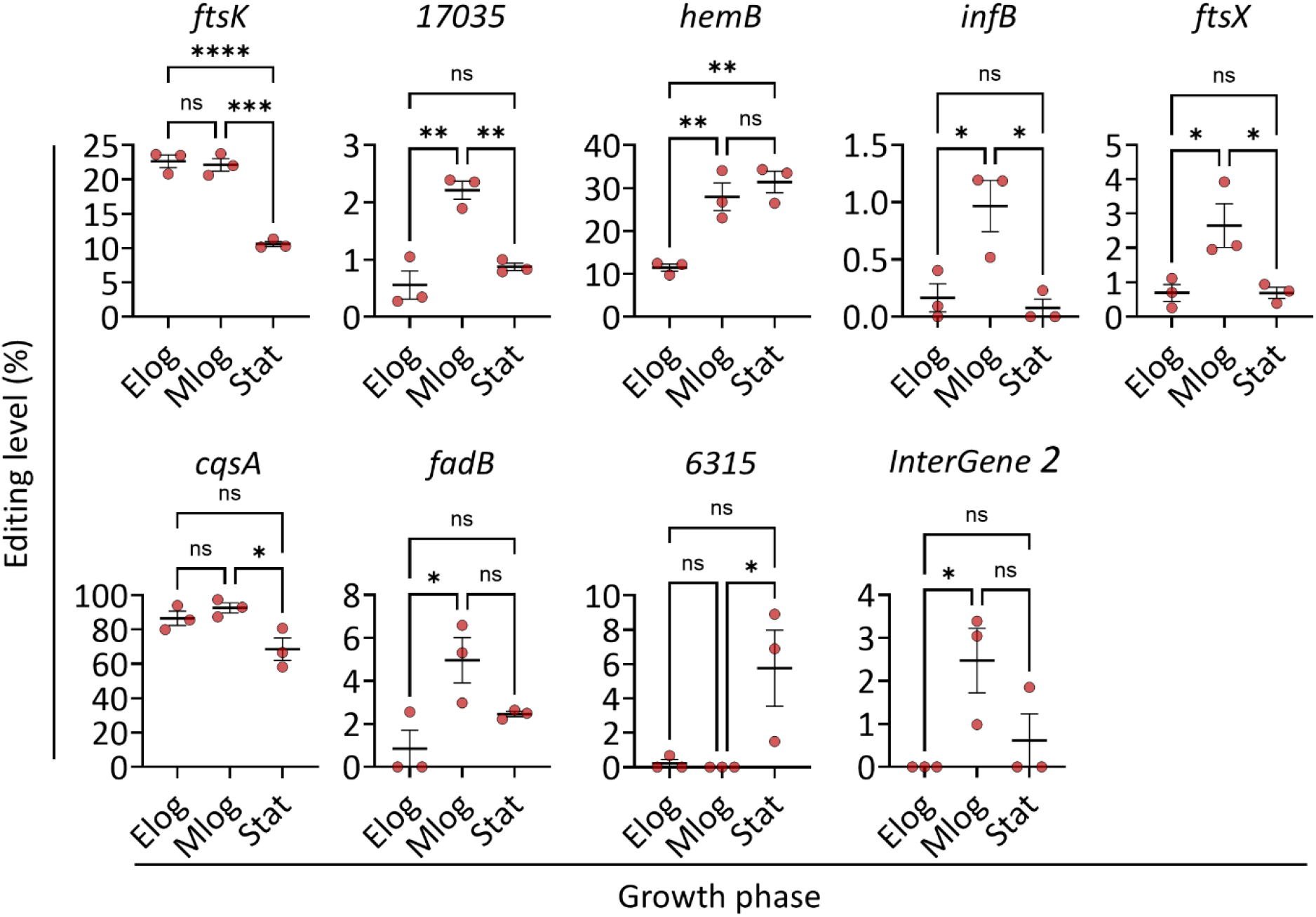
A-to-I mRNA editing in *V. alginolyticus* changes across growth phases in specific transcripts. Editing levels (mean and standard errors) of nine mRNAs showing significant change in editing levels during growth. Statistical analysis was conducted using one-way ANOVA with Tukey’s multiple testing correction. P-value marks are as follows: p ≤ 0.05 (*), p ≤ 0.01 (**), p ≤ 0.001 (***), p ≤ 0.0001 (****); ns – not significant. Elog – Early log phase; Mlog – middle log phase; Stat – Stationary phase.

### A-to-I RNA editing in *V. alginolyticus* occurs in a less conserved sequence motif than in other gammaproteobacteria

In our RNA editing discovery analysis, we focused on sites within a UACG motif (Figure 1) that represents the core sequence required by TadA for editing *in vitro* and *in vivo* (18, 19). However, we recently showed that most of the 381 editing sites found in 64 gammaproteobacterial species (including *Vibrio* species) are embedded within a 7-base motif (YUACGAA) (23). Importantly, about a quarter of the sites were found in a motif that deviates by one base from the 7-base motif, suggesting that different species may have different TadA-recognition motifs.

To examine the sequence context around editing sites in *V. alginolyticus*, we used WebLogo analysis of all 38 UACG RNA editing events, excluding tRNA-Arg2 (Supplementary Table 3). We observed that the motif surrounding editing sites in *V. alginolyticus* was more relaxed compared to our reported gammaproteobacterial motif (23). The most prominent difference was observed in position “-2”, which showed no base preference (Figure 3A). Additionally, position “+4” was less conserved compared to our collective analysis of gammaproteobacterial species (Figure 3A). Thus, although the core UACG motif is conserved across gammaproteobacterial species, the sequences surrounding it can vary between species.

**Figure 3.**
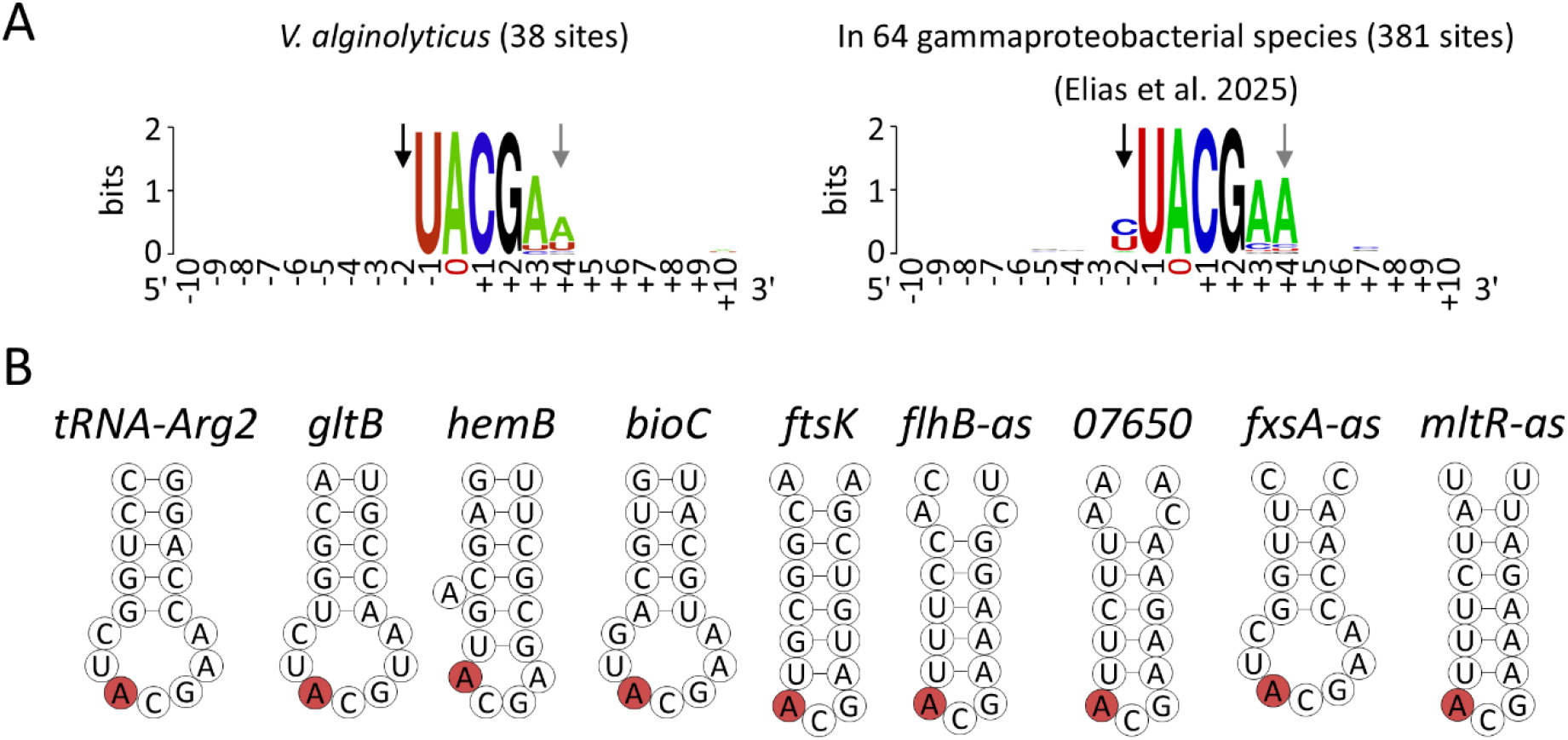
A-to-I RNA editing in *V. alginolyticus* occurs in a shorter conserved sequence motif than other gammaproteobacteria. **A**. WebLogo of the 38 RNA editing events (excluding *tRNA-Arg2*) detected in *V. alginolyticus* compared to the identified motif from 64 gammaproteobacterial species (*including V. alginolyticus*). Position “0” is the edited site. The black and grey arrows designate positions “-2” and “+4”, respectively, in the motif of *V. alginolyticus* and gammaproteobacterial species. **B**. Secondary structure predicted by RNAfold (35) around the A-to-I editing site (red) for the 17 nucleotides composing the anticodon arm of tRNA-Arg2 and the 17 nucleotides around mRNA and antisense RNA edited sites.

Importantly, TadA-dependent tRNA-Arg2 editing requires that the edited site be embedded within a stem-loop structure reminiscent of the anticodon arm of tRNA-Arg2 (19). In addition, we recently showed that most mRNA editing events in gammaproteobacteria are embedded within a stem-loop RNA secondary structure (23). Indeed, analyzing the predicted secondary structure around edited sites in *V. alginolyticus* revealed that most RNA editing events reside in a loop, like the tRNA-Arg2 edited site (26/39; Figure 3 and Supplementary Figure 3).

### A-to-I mRNA editing in *cqsA* is conserved in *Vibrio* species

Among the edited sites, the most edited transcript was *cqsA*. CqsA (Cholerae quorum sensing autoinducer; WP_017635350.1), an enzyme responsible for synthesizing the autoinducer CAI-1 (cholera autoinducer-1) (36, 37). CAI-1 is central in modulating QS pathways, influencing bacterial motility, virulence, and biofilm formation (38, 39). Given the importance of CqsA in bacterial communication, any modifications to its expression or function could have significant implications for *Vibrio* behavior and pathogenicity (38). Thus, we tested the conservation of the editable site in CqsA homologs across *Vibrio* species. We analyzed 102 non-redundant *Vibrio* species genomes and examined the position corresponding to 193 in *V. alginolyticus* CqsA (the site that is recoded by editing; Supplementary Table 5). We observed that a minority of the analyzed *Vibrio* species (3.9%) have a cysteine at position 193 hardcoded in their DNA (Figure 4A). Moreover, many *Vibrio* species (32.4%) from diverse lineages, mainly belonging to the *V. harveyi* and *V. splendidus* clades, encode tyrosine at position 193 (Figure 4A and Supplementary Table 5). Notably, the presence of a tyrosine codon at position 193 suggests that RNA editing may also occur in additional *Vibrio* species, as protein sequences reflect the underlying genetic code and do not account for post-transcriptional modifications, such as editing.

**Figure 4.**
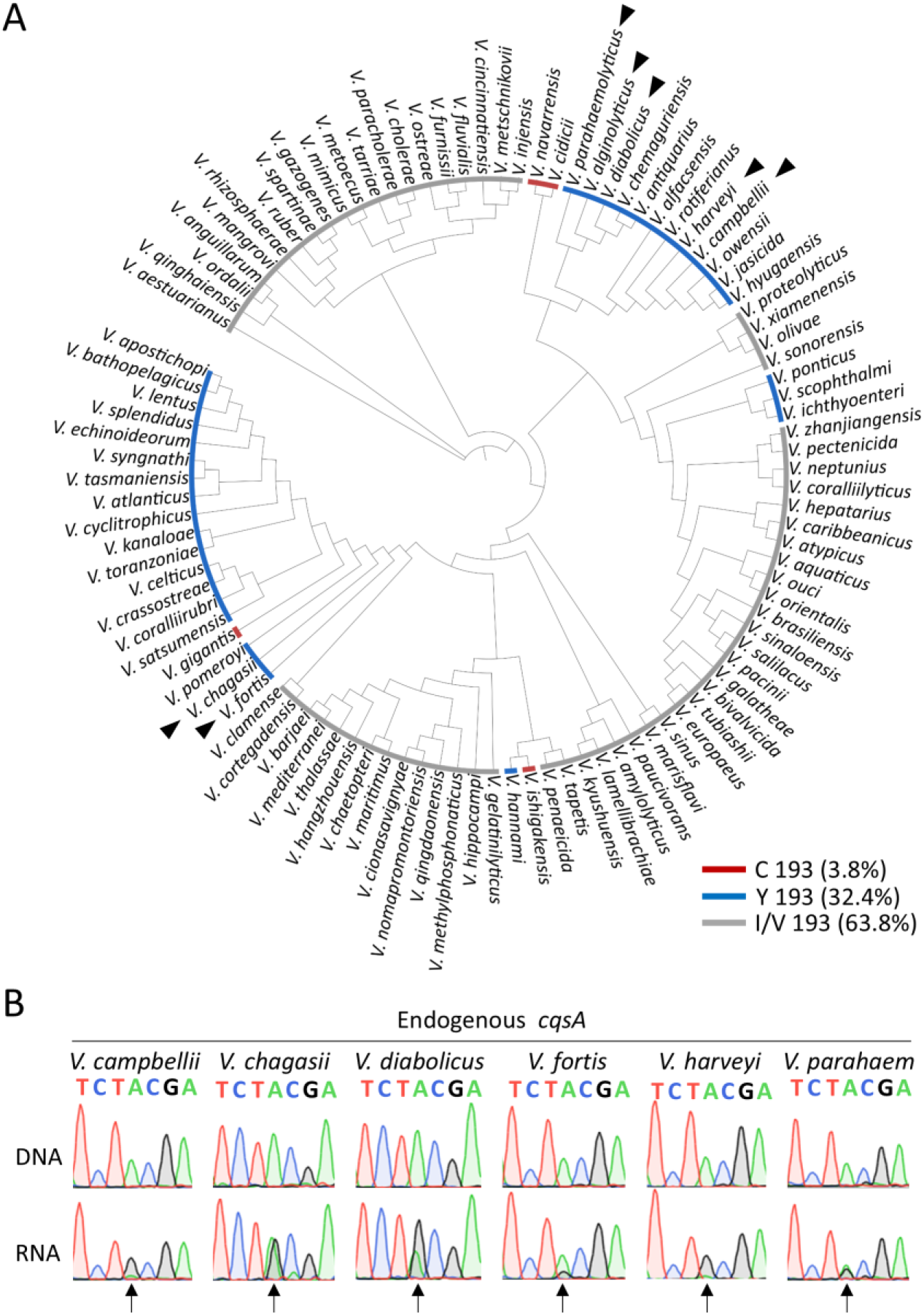
A-to-I mRNA editing in *cqsA* is conserved in multiple *Vibrio* species. **A**. Phylogenetic tree of 102 *Vibrio* species having CqsA (Supplementary Table 5). The tree was constructed using M1CR0B1AL1Z3R based on 1000 core genes shared by all 102 species (40). The identity of the amino acid at position 193 is indicated by blue (Y – non-edited version), red (C – edited version), and grey (I or V – not related to editing) arcs next to the species name. Black triangles mark the species in which RNA editing was validated in *cqsA* in the current study. **B**. Sanger sequencing supports the evolutionary conservation of A-to-I mRNA editing in *cqsA* across six distinct *Vibrio* species, in addition to *V. alginolyticus*. Shown are paired DNA (top) and RNA (cDNA; bottom) chromatograms of each species (extracted from the same sample). The edited site is marked with a black arrow, and the editing signal is represented by a guanosine (black) peak in the RNA samples. The genomic positions of the edited sites are found in Supplementary Table 7. *V. parahaem – Vibrio parahaemolyticus*. All species were grown on MB at 30 °C.

Indeed, we observed that *cqsA* with TadA’s recognition motif was edited in six additional *Vibrio* species, some of which are human pathogens (Figure 4B). In all but one species, the editing level was high, evidenced by the size of the edited base peak (Figure 4B). Importantly, these species represent diverse *Vibrio* lineages (Figure 4A) (26). We conclude that editing in *cqsA* is conserved and likely occurs in additional species, implying a conserved function.

### A-to-I mRNA editing recodes CqsA but does not affect CqsA-mediated quorum sensing

We next examined the effect of editing on CqsA sequence, expression, and QS signaling. To this end, we engineered a strain in which the edited allele is hardcoded at the DNA level, encoding a cysteine at position 193 in the endogenous CqsA (CqsA^EO^; Figure 5A). In parallel, we generated a strain carrying a synonymous tyrosine codon at position 193 that disrupts TadA recognition, thereby preventing editing and producing only the non-edited CqsA isoform (CqsA^NE^; Figure 5A). Notably, whole-genome sequencing confirmed the absence of additional mutations in these strains. Protein mass spectrometry analysis showed that editing recodes the CqsA sequence in the WT strain (Figure 5B and Supplementary Figure 4), whereas the edited peptide was undetectable in the CqsA^NE^ strain (Figure 5B and Supplementary Table 8). Unlike the edited peptide, the non-edited peptide could not be detected with sufficient quality or reliability to meet our high-confidence criteria (Supplementary Table 8; see also Materials and Methods). Finally, total CqsA abundance did not differ significantly between strains, indicating that editing does not affect CqsA expression levels (Figure 5C and Supplementary Table 9).

**Figure 5.**
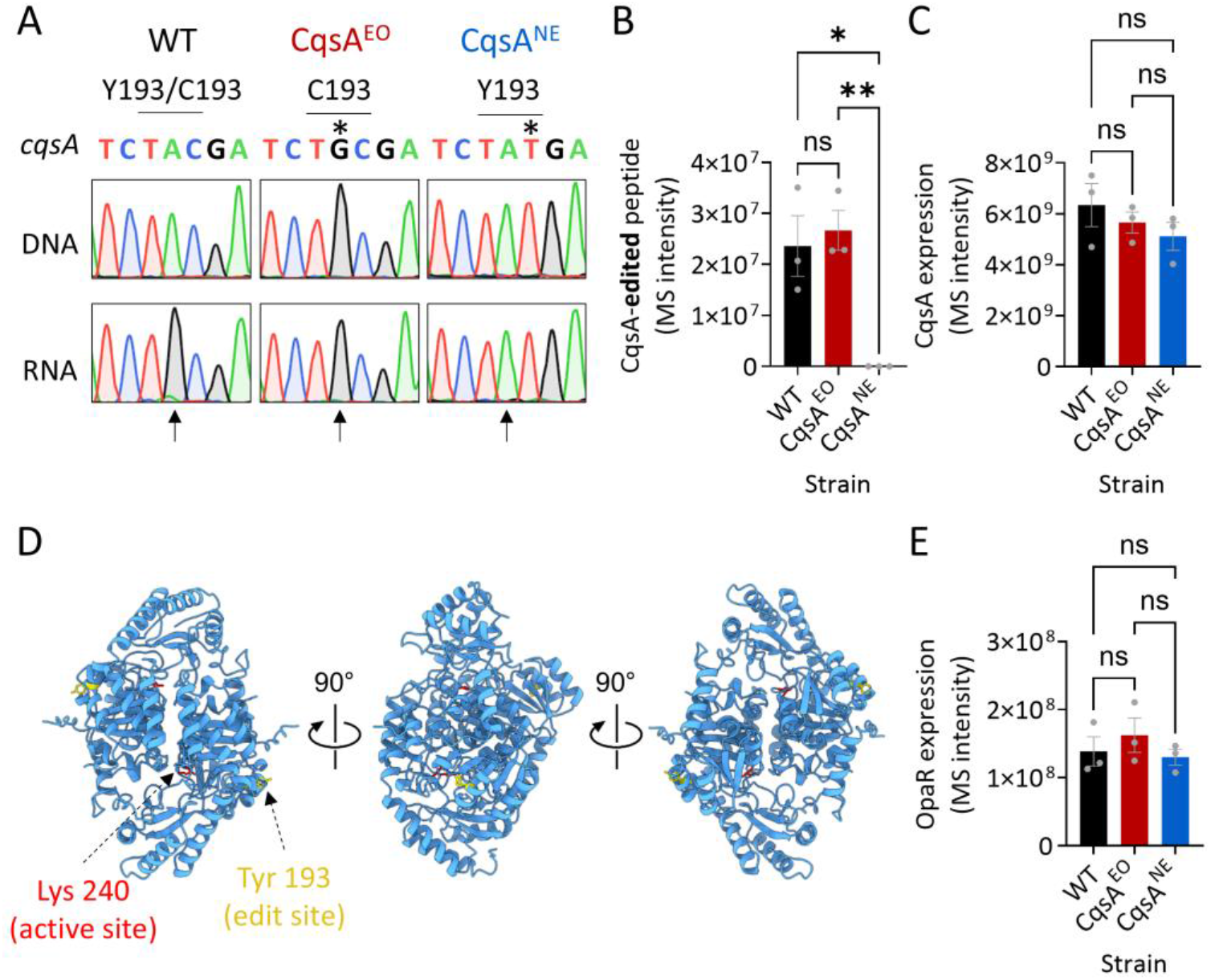
A-to-I mRNA editing recodes CqsA but does not affect CqsA-mediated quorum sensing. **A**. DNA and RNA Sanger sequencing of endogenous *cqsA* in WT and mutant strains of *V. alginolyticus* grown on MB at 30 °C. The mutant strains have a single chromosomal mutation (marked with an *) that results in only the edited (CqsA^EO^; Cys 193) or non-edited (CqsA^NE^; Tyr 193) version of CqsA. **B**. Edited peptide of CqsA abundance (intensity) as detected by protein mass spectrometry in the strains shown in panel A. **C**. CqsA abundance (intensity) as detected by protein mass spectrometry in the strains shown in panel A. **D**. AlphaFold structure prediction of a *V. alginolyticus* CqsA dimer with the edited site (yellow) and active site (red) denoted (43). **E**. Total abundance of OpaR, as detected by protein mass spectrometry in the strains shown in panel A. Statistical analysis in panels B, C, and E was conducted using One-way ANOVA followed by Tukey’s multiple comparisons test: adjusted p-value ≤ 0.05 (*); ≤ 0.01 (**); ns – not significant (p > 0.05).

Structural modeling of a *V. alginolyticus* CqsA dimer, which assumes a similar fold to the known structure of its *V. cholerae* homolog, revealed that the edited residue lies on the surface of CqsA, outside the active site (Figure 5D and Supplementary Figure 5). *Vibrio* have a conserved master QS regulator of the TetR family (e.g., LuxR, HapR, OpaR, and more) that responds to CAI-1 signaling by increasing its own expression and promoting the transcription of hundreds of genes (41, 42). In *V alginolyticus*, OpaR is a LuxR/HapR-type master QS regulator, homologous to *V. harveyi* LuxR (Supplementary Figure 6 and (41)). Consistent with unaltered QS output, OpaR (WP_005379994.1) protein levels were similar in the WT, CqsA^EO^, and CqsA^NE^ strains, suggesting no measurable impact on CAI-1 production or downstream CAI-1-dependent signaling (Figure 5E and Supplementary Table 9). Thus, RNA editing in *cqsA* recodes residue 193 in CqsA without detectably perturbing its canonical QS function, suggesting that editing may instead serve an alternative regulatory role.

### Perturbing A-to-I mRNA editing in *cqsA* affects T6SS-mediated killing

We next examined the effect of RNA editing of *cqsA* on the biology of *V. alginolyticus*. First, we used RNA-seq and differential gene expression analysis to compare gene expression profiles of the WT and mutant strains. We identified 37 and 21 genes with significantly (FDR < 0.01) altered expression in the CqsA^EO^ and CqsA^NE^ strains, respectively, relative to the WT (Figure 6A and Supplementary Table 10). Among these, 17 genes were differentially expressed in the same direction in both mutant strains compared to the WT (Figure 6A and Supplementary Table 10). Thus, perturbing endogenous editing of *cqsA*, either by editing-mimicking (CqsA^EO^) or editing-blocking (CqsA^NE^) mutations, affects gene expression similarly in *V. alginolyticus*.

**Figure 6.**
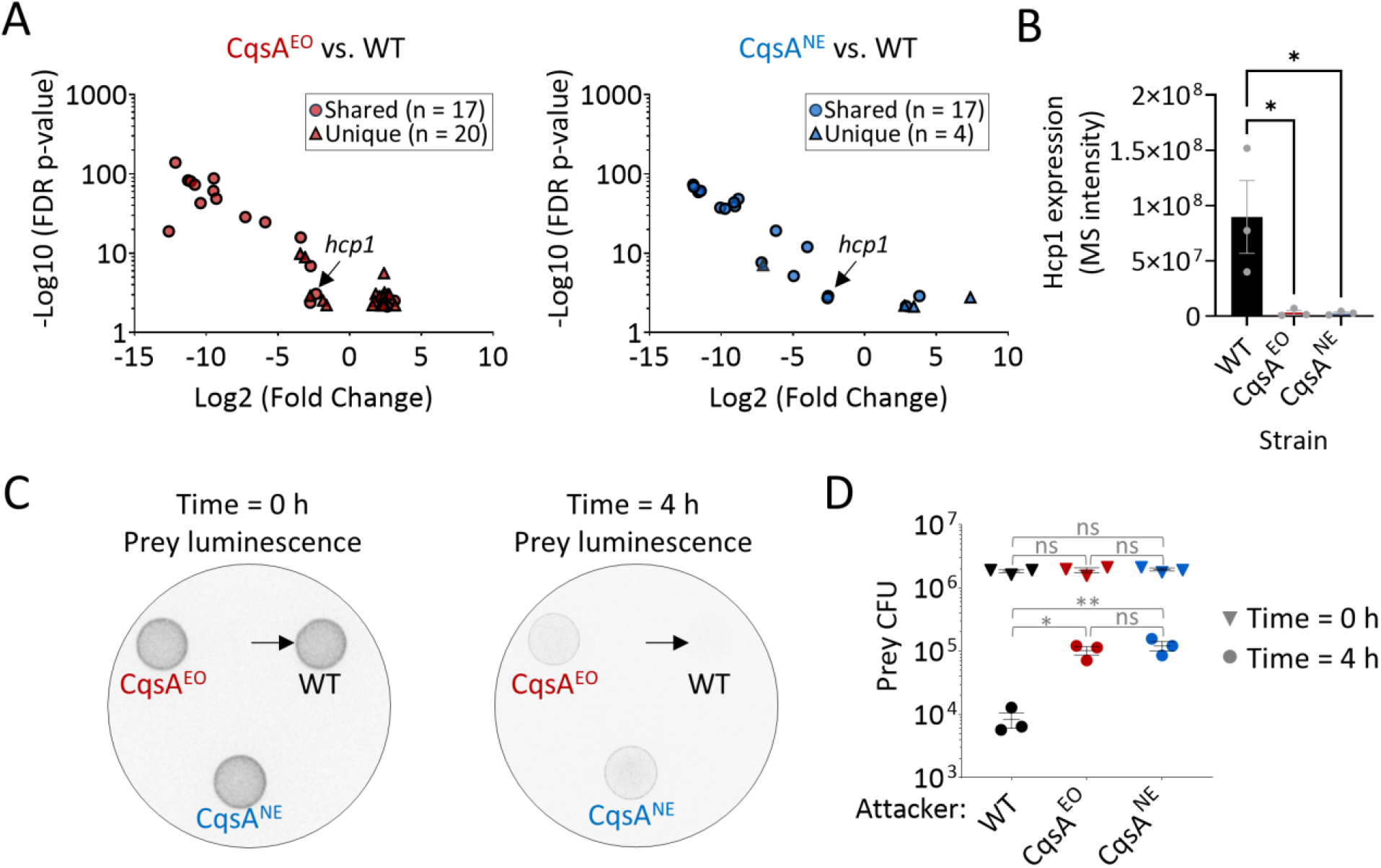
Perturbing A-to-I mRNA editing in *cqsA* affects T6SS-mediated killing. **A**. Volcano plots showing the change in gene expression between the CqsA^EO^ or CqsA^NE^ strains compared to the WT strain at the mid-log phase when grown on Marine Broth at 30 °C. Only genes that are significantly changed are shown (FDR p-value < 0.01). **B**. Total abundance of Hcp1 (WP_005373415.1) as detected by protein mass spectrometry in the CqsA^EO^, CqsA^NE^, and WT strains. **C**. Qualitative assessment of *E. coli* MG1655 prey viability using luminescent signals (prey strains harbor a pCmLux2 plasmid for constitutive expression of the lux operon) before (t = 0 h) and after (t = 4 h) co-incubation with the indicated *V. alginolyticus* attacker strains at a 4:1 (attacker: prey) ratio on MB plates at 30 °C. The experiment was repeated three times, and a representative result is shown. Notice the reduction in the prey luminescent signal when attacked by the WT strain (arrows), indicating efficient killing of the prey. Notably, the presence of all *V. alginolyticus* attacker strains, including the WT, was validated at the same time using white light imaging of the plate (Supplementary Figure 8). **D**. Viability counts (colony-forming units; CFU) of *E. coli* MG1655 prey of the experiment described in panel C. The data are shown as the mean ± SE; n = 3 independent competitions. Statistical analysis in panels B and D was conducted using One-way ANOVA followed by Tukey’s multiple comparisons test: adjusted p-value ≤ 0.05 (*); ≤ 0.01 (**); ns – not significant (p > 0.05).

Among the differentially expressed genes shared between the mutant strains, one that caught our attention was *hcp1*, which was downregulated in both mutant strains compared to WT (Figure 6A and Supplementary Table 10). This gene encodes the core tube component of type VI secretion system 1 (T6SS1), a toxin-delivery apparatus previously shown to inject antibacterial toxins into neighboring bacteria and to mediate interbacterial competition (44). Consistently, we observed a significant reduction in Hcp1 (WP_005373415.1) protein levels in our mass spectrometry data for the mutant strains relative to WT (Figure 6B and Supplementary Table 9). We therefore examined the ability of our strains to intoxicate competing bacteria, using *E. coli* as prey (Supplementary Figure 7). In agreement with the reduced Hcp1 levels, we observed a ten-fold decrease in prey viability following co-incubation with the WT compared to either the CqsA^EO^ or CqsA^NE^ attacker strains (Figure 6C, 6D, Supplementary Figure 8, and Supplementary Table 11). We therefore conclude that editing of *cqsA* affects *V. alginolyticus* killing efficiency, which, to the best of our knowledge, constitutes the first demonstration that endogenous RNA editing has a phenotypic manifestation that affects bacterial competitive fitness.

## Discussion

Previously, by analyzing publicly available RNA-seq data, we reported 14 mRNA editing events in *V. alginolyticus* (23). However, because we had only RNA-seq data available, we used highly stringent parameters to minimize false-positive editing events. As a result, the likelihood of missing real sites increased. Here, we used DNA and RNA sequencing to identify editing events in *V. alginolyticus*. Having both DNA-seq and RNA-seq from the same sample allowed us to exclude signals stemming from DNA variants and sequencing artifacts, and to accurately detect RNA editing events, even at low levels (1% - 5%). As a result, we identified 38 mRNA editing events – three times more editing events than in our previous analysis of *V. alginolyticus* publicly available RNA-seq data (23). Thus, sequencing DNA and RNA from the same sample is superior to analyzing RNA-seq data alone and allows better identification of novel editing events.

In mammals, only a minority of editing events are conserved across species (45). However, among these sites lies the most studied site in the *GRIA2* gene, which alters glutamine to an arginine residue (2). This editing site is essential to the viability of mice. The case of *GRIA2* exemplifies that even a single site can be important for an organism’s fitness and overall well-being. Here, we show that editing of *cqsA* is conserved across *Vibrio* species and has phenotypic consequences, as evidenced by reduced killing activity in the mutant strains (Figure 6C and 6D). Thus, endogenous editing events can affect bacterial behavior, as was shown in metazoans (15, 46-48).

CqsA is well known as the auto-inducer synthase that controls QS. However, we did not detect an effect on QS signaling, as evidenced by OpaR protein expression levels (Figure 5E), when we mutated the edited site. In addition, we observed only 17 genes that were differentially expressed in the same direction in both mutant strains. This is in sharp contrast to RNA-seq studies in *V. harveyi* in which perturbation of the QS pathway affects hundreds of genes. (42). Moreover, the edited site (Y193C) is located on the surface of CqsA, not near its catalytic site (Figure 5D). So, it is possible that CqsA and editing within have other functions unrelated to QS and auto-inducer synthesis. Our data indicates that CqsA is edited to an average of 70-90% depending on the growth phase (Figure 2). Thus, in theory, there are 10-30% non-edited CqsA as well. It could be that the role of editing is to create two CqsA isoforms, whose equilibrium fine-tunes the regulation of a subset of genes, including the T6SS1 component *hcp1*, thereby affecting competitive fitness. This scenario is supported by the fact that both the editing-mimicking (CqsA^EO^) and editing-blocking (CqsA^NE^) strains exhibit similar gene expression patterns and killing efficiency in *V. alginolyticus*. Furthermore, CqsA was shown to affect the expression and activity of T6SS in other *Vibrio* species, further supporting a role for CqsA editing in T6SS regulation (49, 50). Thus, our results, combined with the known effect of CqsA on T6SS in other *Vibrio* species, suggest that having only a single CqsA variant, either edited or non-edited, is insufficient. It is also noteworthy that we observed additional genes that were differentially expressed in the mutants compared to the WT, suggesting broader effects of editing in *V. alginolyticus*. However, the molecular mechanisms underlying CqsA editing–related phenotypes in *V. alginolyticus* remain to be elucidated.

Finally, we identified A-to-I RNA editing events in 37 additional genes/transcripts beyond *cqsA* (Figure 1). The conservation and functional importance of the edited site in *cqsA* suggest that other sites may also be conserved and carry yet-to-be-discovered functions across *Vibrio* species. Thus, future studies should systematically examine the landscape of A-to-I RNA editing across *Vibrio* species and test its functional roles.

Our research identifies *V. alginolyticus* as the bacterial species with the highest number of confirmed A-to-I RNA editing events and, for the first time, shows that both non-edited and edited “RNA alleles” of a single gene are required. Thus, A-to-I RNA editing of a coding sequence in its natural context can have functional consequences for bacterial behavior and fitness.

## Acknowledgments

We thank the European Research Council (Project 101116636 — REDBAC to D. B.-Y.) for supporting this research. D.S. was supported by the Israel Science Foundation (ISF grant number 1362/21). C.M.F was supported by a Gray Scholarship for Post-Doctoral Fellows, Gray Faculty of Medical and Health Sciences Award, Tel Aviv University. We thank Ben-Gurion University of the Negev genomic core for producing the RNA-seq DNA-seq data. We thank the Smoler Proteomics Center at the Technion for the protein mass spectrometry analysis.

## Data Availability Statement

The DNA- and RNA-seq data were deposited to the NCBI SRA under accession PRJNA1320707 and will become publicly available upon publication.

The mass spectrometry data were deposited to Pride through ProteomeXchange in Project accession: PXD078381

## Materials and Methods

### Bacterial strains

During the course of this work, we used the following strains and growth conditions: *V. alginolyticus* ATCC 17749 was grown on Marine Broth (MB) media (Millipore #76448) at 30 °C or 37 °C (depending on the downstream analysis); *E. coli* DH5α (λ-Pir) and *E. coli* HB101 containing the pRK2013 plasmid were grown in Lysogeny broth (LB) at 37 °C; *Vibrio harveyi* TBV014, *Vibrio fortis* TBV008, *Vibrio chagasii* TBV018, and *Vibrio diabolicus* TBV026, isolated from Israel coastal waters (51), as well as *Vibrio campbellii* ATCC BAA-1116 and *Vibrio parahaemolyticus* RIMD 2210633, were grown on MB media (Millipore #76448) at 30 °C.

### DNA and RNA extractions, and cDNA synthesis

At mid-log (OD_600_ of 0.5 – 1) or stationary phase (24 hours of growth), 1.5 mL were taken for DNA and RNA extractions. DNA was extracted using a GeneJET Genomic DNA Purification Kit (Thermo Scientific, #K0721) or the iNtRON biotechnology G-spin™ Genomic DNA Purification Kit (Life gene # 17121), while RNA was extracted using the GeneJET RNA Purification Kit (Thermo Scientific #K0731). RNA samples were treated with 4 units of DNase I (NEB # M0303L) for 20 minutes at 37 °C. Finally, following incubation of 15 minutes at 65 °C, cDNA synthesis was done using GoScript Reverse Transcription Mix (Promega # A2801). To synthesize cDNA, 500 nanograms of total RNA were primed with random hexamers and reverse-transcribed with the GoScript Reverse Transcription Mix kit (A2801 Promega) following the manufacturer’s protocol.

### RNA editing discovery in *V. alginolyticus*

Each sample was derived from a single and different colony and was grown in MB medium. RNA was extracted at early lag (OD_600_ of 0.15-0.24), mid-log (OD_600_ of 1.3-1.6), and stationary phase (24 hours of growth; OD_600_ above 3.3), as described above. Ribosomal RNA was depleted using NEBNext® rRNA Depletion Kit (Bacteria) (New England Biolabs #E7850). RNA-seq Libraries were constructed using NEBNext® Ultra™ II Directional RNA Library Prep Kit for Illumina® (New England Biolabs, #E7760). DNA-seq libraries were prepared using NEBNext® Ultra™ II FS DNA Library Prep Kit for Illumina (New England Biolabs, #E7805L). Finally, RNA-seq and DNA-seq libraries were sequenced on the NovaSeq X platform (Illumina).

We used CLC Genomics Workbench for all steps of the analysis (described below).

RNA-seq reads were first trimmed according to length and quality scores to ensure high quality of the reads by using the following parameters: Trim using quality scores = Yes; Quality limit = 0.01; Trim ambiguous nucleotides = Yes; Maximum number of ambiguities = 1; Automatic read-through adapter trimming = Yes; Minimum length = 50; Maximum length = 150; Remove 5’ terminal nucleotides = No; Remove 3’ terminal nucleotides = No; Remove on first read = Yes; Remove on second read (for paired reads) = Yes; Trim to a fixed length = No; Trim end = Trim from 3’-end; Discard short reads = Yes; Discard long reads = No; Save discarded sequences = No; Save broken pairs = No.

Paired RNA reads were merged into a single longer read maintaining the original orientation of the R1 reads (reverse to the original transcript direction). The parameters were as follows: Mismatch cost = 2; Minimum score = 8; Gap cost = 3; Maximum unaligned end mismatches = 0.

Next, RNA-seq and DNA-seq reads were mapped to the *V. alginolyticus* reference genome (NC_022349.1 and NC_022359.1) with the following parameters: Masking mode = No masking; Match score = 1; Mismatch cost = 2; Cost of insertions and deletions = Linear gap cost; Insertion cost = 3; Deletion cost = 3; Length fraction = 0.95; Similarity fraction = 0.95; Global alignment = No; Non-specific match handling = Ignore.

Following the mapping step, initial RNA variant calling was performed using the following parameters: Ignore positions with coverage above = 100,000; Restrict calling to target regions = Not set; Ignore broken pairs = Yes; Ignore non-specific matches = Reads; Minimum coverage = 4; Minimum count = 2; Minimum frequency (%) = 0.1; Base quality filter = Yes; Neighborhood radius = 5; Minimum central quality = 30; Minimum neighborhood quality = 30; Read direction filter = No; Relative read direction filter = No; Read position filter = No.

To exclude DNA variants, RNA variants were filtered against the DNA samples. Variants with more than one supporting read at the DNA-seq dataset were removed.

After the initial variant calling was performed, additional filtering was applied to obtain high-confident variants, with sufficient coverage (>=3 reads) and frequency (>=1%). All filtered variants were required to match all the following criterions: Criteria = ‘Type contains SNV’; Criteria = ‘Reference allele contains No’; Criteria = ‘Frequency >= 1’; Criteria = ‘# unique start positions >= 3’; Criteria = ‘# unique end positions >= 3’; Criteria = ‘read Count >= 3’; Criteria = ‘Read coverage >= 10’; Criteria = ‘Reference contains A and the variant contains G’ or ‘Reference contains T and the variant contains C.

Next, we filtered variants shared between at least two (out of three) biological replicates of each growth phase. Thus, to be identified as an editing event, a variant was required to be present in a minimum of two out of three samples and comprise at least 1% of the reads, alongside the additional parameters described above.

Finally, we extracted the status of the identified shared variants from the previous step from the RNA mapping step. We did this to make sure we did not missed variants that were filtered out due to our initial parameters.

### RNA editing motif analysis

Using CLC Genomics Workbench, we extracted the four-base sequences surrounding all A-to-G mismatches. The expected frequencies of each four-base motif were calculated and compared to the observed motif frequencies using a chi-square test for goodness of fit.

TadA’s sequence motif was identified using WebLogo analysis (52). In short, we extracted the 21 bases surrounding each mRNA edited site (10 bases upstream and downstream) and analyzed them using the WebLogo Server with default parameters.

### Secondary structure prediction around editing events

We used RNAfold from the ViennaRNA Package 2.0 (35) to calculate the minimum free energy (MFE) around editing events. As previously shown on editing events in *S. pyogenes* and gammaproteobacterial species (20, 23), we calculated the structure of the minimum free energy of 17 nucleotides around edited sites (which is the length of tRNA-Arg2 anticodon). We position the edited adenosine as the ‘0’ position in our sliding window analysis similarly to its location in the anticodon arm of *tRNA-Arg2*.

### RNA editing evolutionary conservation analysis

In order to gain an evolutionary perspective on the effect of editing within *cqsA*, we aimed to characterize the amino acid identity across *Vibrio* species. We used CqsA protein sequence of *V. alginolyticus* as a query in BlastP analysis against up to 5000 *Vibrio* species targets (the maximum allowed number of targets in NCBI’s web-blast) in the RefSeq database, with an e-value of E-05 (53). BlastP identified 2669 CqsA homologs. We used the MAFFT server for multiple sequence alignment and MEGA to extract the amino acid at the edited site (54, 55). Finally, redundant sequences, partial sequences, missing data, or sequences of uncultured/unclassified species, as well as unnamed or hypothetical proteins were excluded, resulting in 105 CqsA sequences from 102 species. Notably, three species had two versions of CqsA, each with a different amino acid at position 193 (*Vibrio atypicus, Vibrio scophthalmi*, and *Vibrio sinaloensis*).

To put the identity of amino acid at position 193 in an evolutionary context, we download the genomes of the 102 species having CqsA (Supplementary Table 6) and used M1CR0B1AL1Z3R (40) to construct a phylogenetic tree. M1CR0B1AL1Z3R identified 1062 core genes and selected 1000 for the construction of the tree according to its default parameters.

### Mutating *cqsA* in *V. alginolyticus*

To mutate the editing motif in *cqsA* in *V. alginolyticus*, 1 kb sequences upstream and downstream of the edited site in *cqsA* gene were cloned into pDM4, a Cm^R^OriR6K suicide plasmid (56). Next, we created two plasmid versions by mutating the edited site (A-to-G; recoding a tyrosine to a cysteine codon; “edited” CqsA plasmid) or the adjacent downstream base (C-to-T; mutating one tyrosine codon to another; “non-edited” CqsA plasmid). To create the edited and non-edited plasmids, we amplify the entire plasmid using high-fidelity Kapa DNA polymerase (Roche) with back-to-back primer pairs containing a single-point mutation at a desired location. Next, products were phosphorylated with T4 Polynucleotide Kinase (NEB), ligated with T4 DNA ligase (NEB), and treated with Dpn1 (NEB) to eliminate any residual plasmid DNA used as PCR template.

The pDM4 constructs were transformed into *E. coli* DH5α (λ-Pir) by heat shock and then transferred into *V. alginolyticus* via conjugation, as previously described (57). Transconjugants were selected on agar plates supplemented with chloramphenicol (35 µg/mL), and then counter-selected on agar plates containing 15% (wt/vol) sucrose for loss of the *sacB*-containing plasmid. Through this methodology, we successfully generated two *V. alginolyticus* strains: non-edited *cqsA* strain (NC_022359; G1173397A; Tyr193Tyr) and a mutant strain mimicking 100% editing in *cqsA* (NC_022359; T1173398C; Tyr193Cys). Mutations were confirmed by PCR and Sanger sequencing. Primers for all PCR reactions are provided in Supplementary Table 7. The sequences were aligned and visualized using SnapGene (Dotmatics).

### Mass spectrometry

#### Proteolysis

Bacterial cells were lysed in 8.5 M Urea, 400 mM ammonium bicarbonate, and 10 mM DTT, sonicated twice (90%, 10-10, 5’), and centrifuged (10,000 g, 10’). Protein concentration was estimated using the Bradford method. The samples were reduced (60°C for 30 minutes), modified with 35.2 mM iodoacetamide in 100 mM ammonium bicarbonate (room temperature for 30 minutes in the dark), and digested in 1 M Urea and 66 mM ammonium bicarbonate with chymotrypsin (V106A, Promega) overnight at 25°C in a 1:50 (M/M) enzyme-to-substrate ratio. An additional digestion step with chymotrypsin was done for 4 hours at 25°C in a 1:100 (M/M) enzyme-to-substrate ratio.

Mass spectrometry analysis: The resulting peptides were analyzed by LC-MS/MS using an Exploris 480 mass spectrometer (Thermo) fitted with a capillary HPLC (EV-1000, Evosep One). The peptides were loaded onto a 15 cm, ID 150 µm, 1.9-micron Performance column EV1137 (Evosep). The peptides were eluted with the built-in Xcalibur 15 SPD (88 minuntes) method. Mass spectrometry was performed in a positive mode (m/z 280-1500, resolution 120,000 for MS1 and 15,000 for MS2) using 2 scan events. In the first, a repetitive full MS scan was followed by high collision dissociation (HCD, at 27 normalized collision energy) of the 20 most dominant ions (charges 2-6) selected from the first MS scan. The AGC settings were 3×106 for the full MS and 1×105 with maximum injection time of 30 ms for the MS/MS scans. The intensity threshold for triggering MS/MS analysis was 8×103. A dynamic exclusion list was enabled with an exclusion duration of 30 s. In the second scan event, repetitive full MS scan (m/z 280–1500) was followed by high collision dissociation (HCD, at 27 normalized collision energy) of the 12 most dominant ions (charges 2-6) selected from a specific peptide mass list. The AGC setting for MS/MS was 1×105 with maximum injection time of 100 ms. An exclusion list was not enabled in this scan event.

#### Data analysis

The mass spectrometry data were analyzed with Proteome Discoverer 2.4 (Thermo) using Sequest search engine, searching against the *Vibrio alginolyticus* (Feb 2026, 4596 entries) proteome from the Uniprot database and the specific sequence of CqsA (A0A0L7TKN2; edited and non-edited), with mass tolerance of 20 ppm for the precursor masses and 0.02 Da for the fragment ions. Oxidation on methionine and protein N-terminus acetylation were accepted as variable modifications, and carbamidomethyl on cysteine was accepted as a static modification. The minimal peptide length was set to six amino acids, and a maximum of two miscleavages was allowed. The data were quantified by label-free analysis using the same software. Peptide-level false discovery rates (FDRs) were filtered to 1% using the target-decoy strategy.

Spectra that were not confidently identified in the initial search were re-searched against a reduced database containing only the edited and non-edited CqsA sequences, allowing semi-chymotryptic peptides to detect potential non-specific cleavages present in the samples. Because of the restricted search space and relaxed cleavage specificity, peptide identifications obtained from this secondary search were treated as lower confidence. For downstream analysis, a peptide was considered high confidence only if it met all of the following criteria: (i) it was identified in the initial database search (indicating a reliable peptide backbone fragmentation pattern in MS/MS); (ii) it was consistent with complete chymotryptic digestion; (iii) it exhibited the expected strain specificity, such that the edited peptide was absent from the CqsA^NE^ strain and the non-edited peptide was absent from the CqsA^EO^ strain; and (iv) it was independently identified in at least 2 out of 3 biological replicates. Only the edited peptide fulfilled these criteria, whereas the non-edited peptide did not (Supplementary Table 8).

### Structural modeling

The structure prediction of the *V. alginolyticus* CqsA (WP_017635350.1) dimer was carried out using AlphaFold 3 (58) (https://alphafoldserver.com/). The best model was selected and visualized using ChimeraX (59) version 1.7.1. The solved structure of the *V. cholerae* CqsA dimer was downloaded from PDB (3kki) and superimposed onto the predicted structure of the *V. alginolyticus* CqsA prediction.

### Differential gene expression analysis

Bacterial culture preparation: *V. alginolyticus* strains (WT, CqsA^EO^, and CqsA^NE^) were grown in triplicate in MB media, with each biological replicate initiated on a different day from a single colony. At mid-log phase (OD_600_ = 0.61-0.92), 1 mL of culture was collected, and the cells were harvested by centrifugation for RNA extraction.

RNA extraction: The supernatants were discarded, and RNA was extracted from the bacterial pellets using the GeneJET RNA Purification Kit (Thermo Scientific #K0731). RNA samples were treated with 4 units of DNase I (NEB # M0303L) for 20 minutes at 37 °C. RNA integrity was determined on a QIAxcel device (QIAGEN, QIAxcel RNA QC Kit v2.0), and concentration was determined using a QuantiFluor® RNA System (Promega, #E3310) on a Qubit™ Flex Fluorometer (Invitrogen).

RNA-seq analyses: RNA-seq libraries were constructed using the Zymo-Seq RiboFree Total RNA Library Kit per manufacturer’s instructions (Zymo Research, #R3003). The molarity of libraries was determined using QIAxcel (QIAxcel DNA High Sensitivity Kit) and QuantiFluor® dsDNA System (Promega, # E2670) on a Qubit™ Flex Fluorometer (Invitrogen). Libraries were sequenced (150PE) on a NovaSeq X (Illumina).

CLC Genomics Workbench was used for all analysis steps as described follows: RNA-seq reads were first trimmed according to length and quality scores to ensure high quality of the reads by using the following parameters: Trim using quality scores = Yes; Quality limit = 0.01; Trim ambiguous nucleotides = Yes; Maximum number of ambiguities = 1; Automatic read-through adapter trimming = Yes; Trim homopolymers from 5’ = No; Trim homopolymers from 3’ = Yes; polyA = No; polyC = No; polyG = Yes; polyT = No; Remove 5’ terminal nucleotides = No; Remove 3’ terminal nucleotides = No; Remove on first read = Yes; Remove on second read (for paired reads) = Yes; Trim to a fixed length = No; Maximum length = 150; Trim end = Trim from 3’-end; Discard short reads = Yes; Minimum length = 50; Discard long reads = No; Save discarded sequences = No; Save broken pairs = No.

Next, we performed differential gene expression analysis. To obtain normalized expression measurements (as transcripts per million, TPM), we mapped the RNA-seq reads (following quality control and trimming of adapter read-through) to the Genome of *V. alginolyticus* ATCC 17749 (NCBI RefSeq NC_022349.1 and NC_022359.1) with the following parameters in CLC Genomics Workbench: Reference type = Genome annotated with genes only; Reference sequence = *V. alginolyticus* ATCC 17749; Gene track = Vibrio alginolyticus ATCC 17749_Gene; Use spike-in controls = no; Mismatch cost = 2; Insertion cost = 3; Deletion cost = 3; Length fraction = 0.5; Similarity fraction = 0.8; Global alignment = No; Strand specific = Reverse; Library type = Bulk; Maximum number of hits for a read = 10; Count paired reads as two = No; Ignore broken pairs = No; Expression value = TPM. Finally, we analyzed differential gene expressions using default parameters (also using CLC Genomics Workbench).

### Interbacterial competition assays

The competition assays were performed as previously described, with minor modifications (60). Briefly, to evaluate interbacterial competition using CFU counts, attacker and prey strains, the latter containing the pCmLux2 plasmid ((61); providing a selective marker), were grown overnight in appropriate media in triplicate. Bacterial cultures were normalized to an OD_600_ of 0.5 and mixed at a 4:1 (attacker:prey) ratio. Bacterial mixtures were spotted onto MB agar plates (25 µL per spot) and incubated at 30 °C. To evaluate prey viability at the beginning of the experiment (t = 0 h), 25 µL were collected from all competition mixtures immediately after mixing and added to 475 µL of MB media. Then, the samples were serially diluted (10-fold) and plated on selective LB agar plates supplemented with chloramphenicol (10 µg/mL) to allow growth of the prey strain. Prey CFUs were counted after an overnight incubation. To evaluate prey viability at the end of the experiment (t = 4 h), each competition spot was scraped into 500 µL of MB media, serially diluted (10-fold), and plated as described above. Plates were incubated overnight, and colonies were counted the following day. To evaluate prey viability via luminescence, agar plates were visualized at t = 0 h and t = 4 h in a Fusion FX6 imager (Vilber Lourmat) using a luminescence setup (luminescence was produced by the *lux* operon, constitutively expressed in the *E. coli* prey strain from the pCmLux2 plasmid). The assays were performed three times with similar results.

### Statistical analysis

We used PRISM 10 to conduct the statistical analyses described in the text.

## Supplementary Figures

**Supplementary Figure 1.**
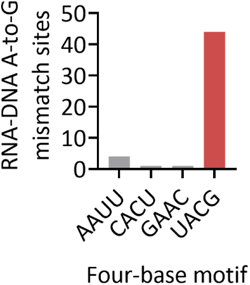
Sequence motif distribution around all 50 A-to-G RNA-DNA mismatches in their genomic context identified in *V. alginolyticus*.

**Supplementary Figure 2.**
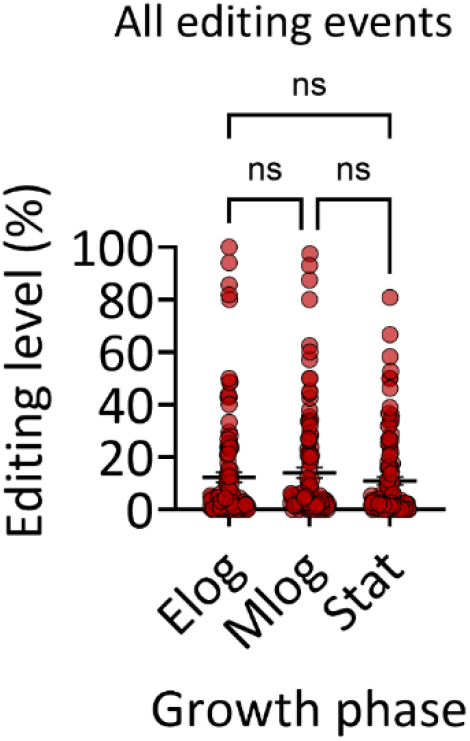
Average and standard errors of editing levels across 38 edited mRNAs as measured across growth phases in MB. Statistical analysis was conducted using one-way ANOVA to examine differences between growth phases, followed by Tukey’s multiple-comparison correction (ns – not significant).

**Supplementary Figure 3.**
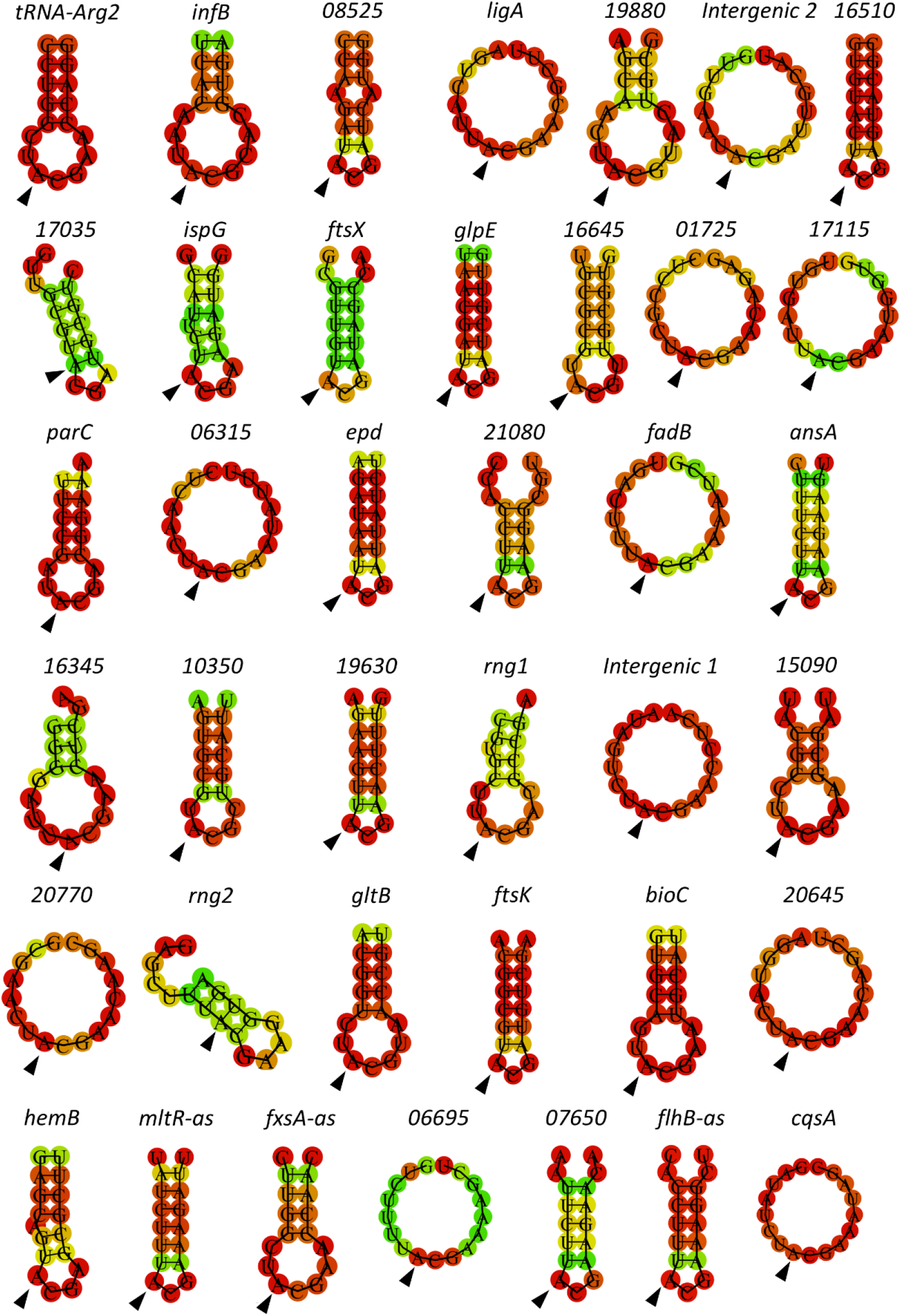
Minimum free energy (MFE) secondary structure predicted by RNAfold (35) around A-to-I editing sites (marked by a black arrowhead). Predictions were made using 17 nucleotides around the edited site in all 38 edited mRNAs and in tRNA-Arg2 of *V. alginolyticus*, as the original output from RNAfold.

**Supplementary Figure 4.**
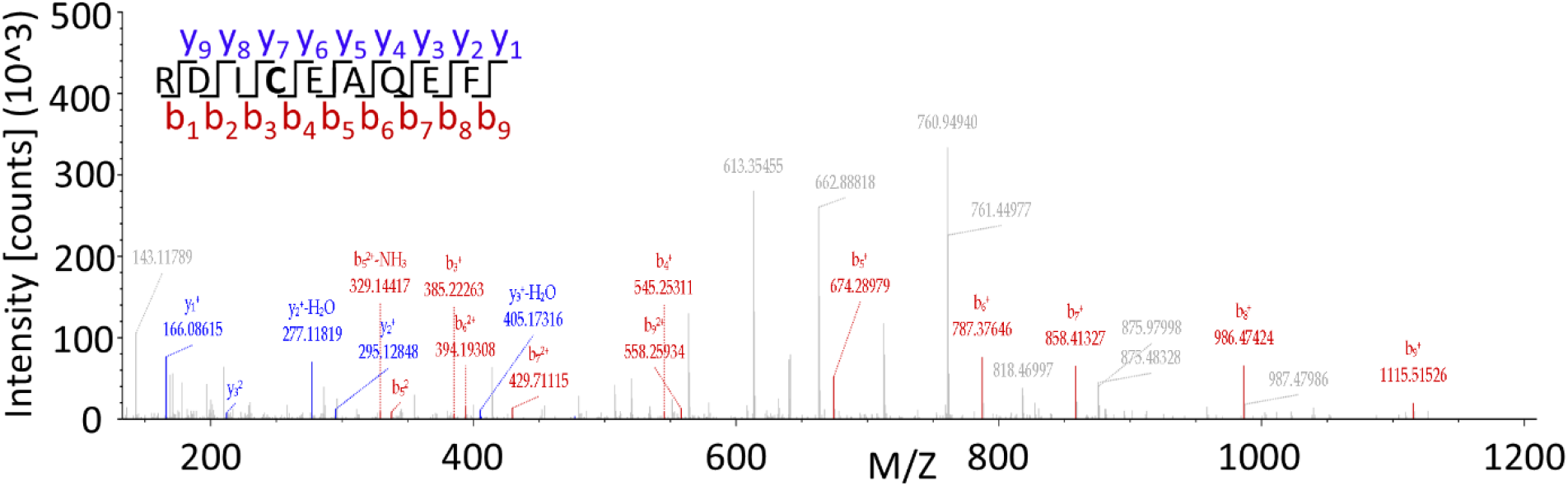
A-to-I mRNA editing recodes the CqsA protein sequence. Protein mass spectrometry spectra of the edited peptide of CqsA as discovered in the WT strain in all 3 replicates) (Supplementary Table 8).

**Supplementary Figure 5.**
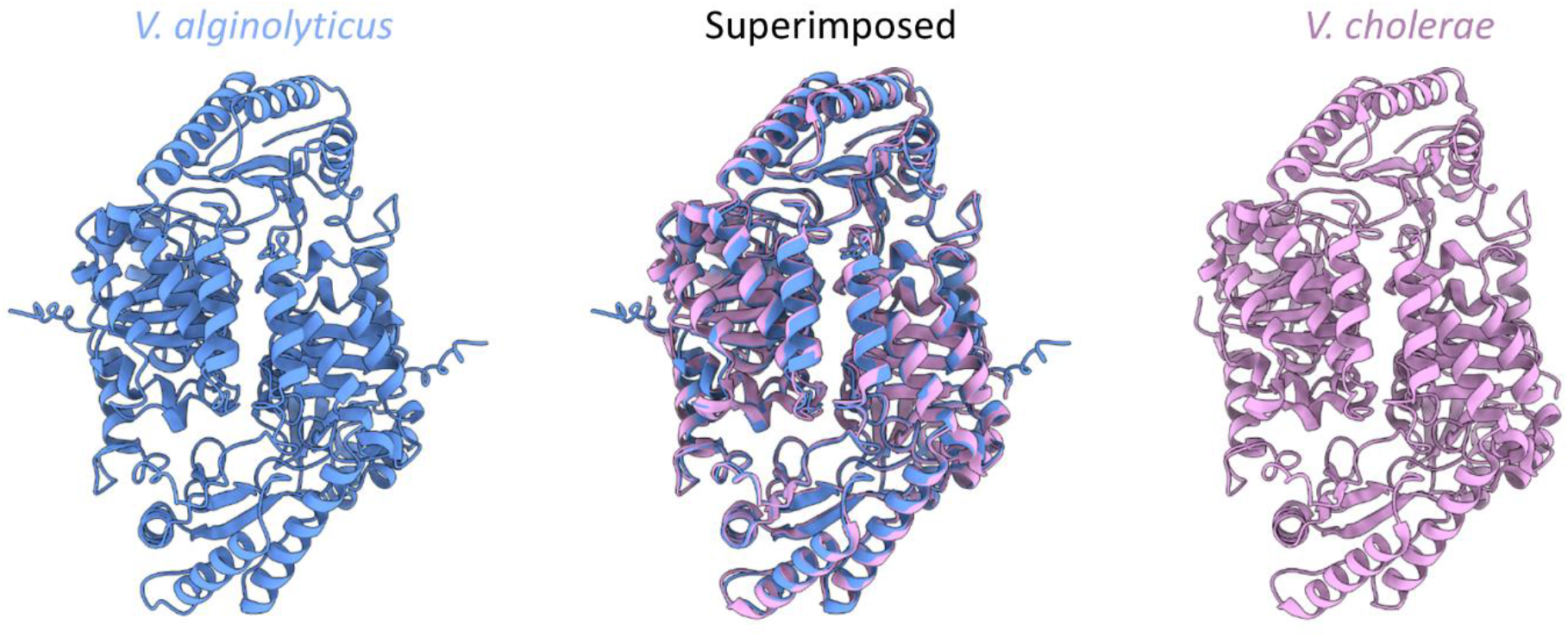
Comparison between the AlphaFold-predicted CqsA dimer from *V. alginolyticus* and the solved CqsA dimer structure from *V. cholerae*. Shown are the AlphaFold 3-predicted structure of *V. alginolyticus* (blue), the solved structure of *V. cholerae* (pink; PDB: 3kki), and their superimposition illustrating the high structural similarity between the two dimers (RMSD = 0.733 Angstroms).

**Supplementary Figure 6.**
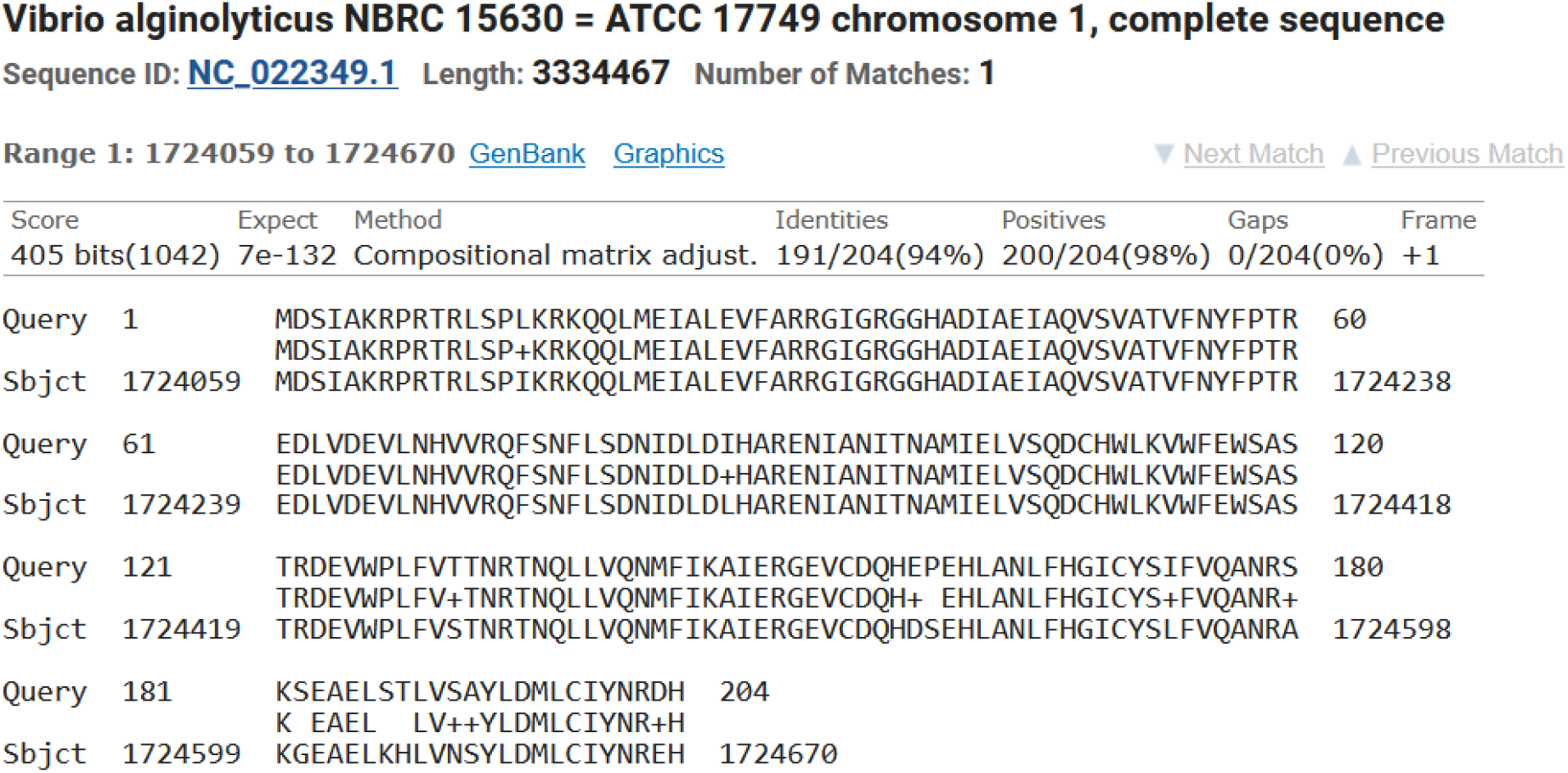
OpaR is a LuxR homolog in *V. alginolyticus*. tBLASTn analysis (62) was performed using the *V. harveyi* LuxR protein sequence (GenBank: SQA34038.1) as the query against both chromosomes of the *V. alginolyticus* genome (NCBI reference sequences NC_022349.1 and NC_022359.1). The identified protein shares 94% amino acid identity with LuxR and is annotated as OpaR on chromosome 1 (NC_022349.1). Shown are the coordinates of the protein (query) and genome (subject) sequences.

**Supplementary Figure 7.**
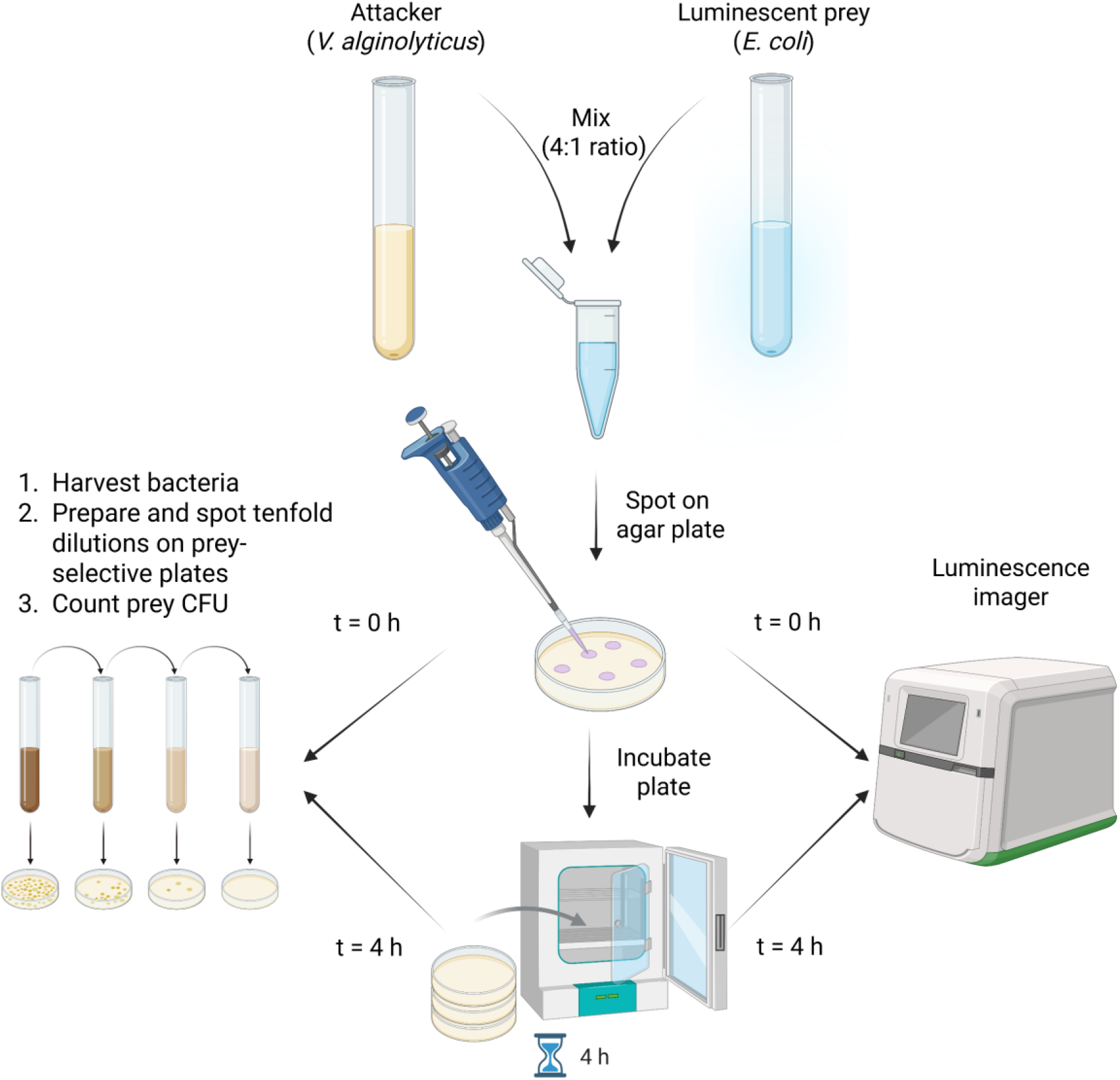
Interbacterial competition assays (related to Figure 6C and 6D). The image was compiled using Biorender.com.

**Supplementary Figure 8.**
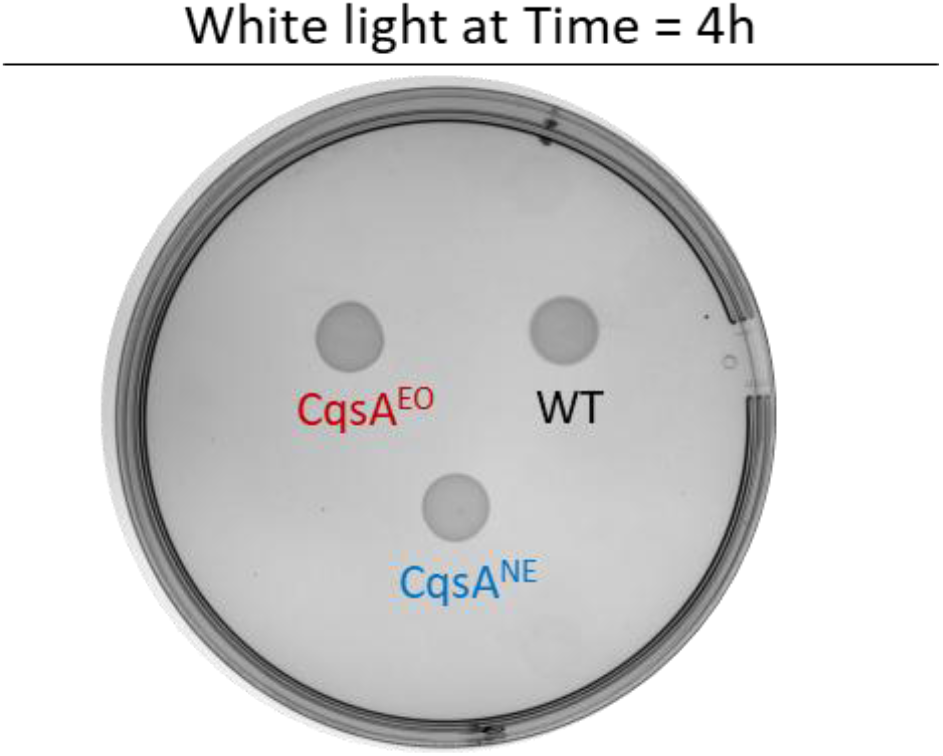
Supporting image for Figure 6C. White light imaging validates the presence of the *V. alginolyticus* attacker strains after co-incubation at a 4:1 (attacker: prey) ratio on MB plates at 30 °C. The experiment was repeated three times, and a representative result is shown. Notice the presence of the WT attacker strain, which is not visible in Figure 6C, where we measure only luminescence originating from the prey strain.

## References

1. E. Eisenberg, E. Y. Levanon, A-to-I RNA editing—immune protector and transcriptome diversifier. Nature Reviews Genetics 19, 473–490 (2018).

2. B. Sommer, M. Köhler, R. Sprengel, P. H. Seeburg, RNA editing in brain controls a determinant of ion flow in glutamate-gated channels. Cell 67, 11–19 (1991).

3. M. Higuchi et al., Point mutation in an AMPA receptor gene rescues lethality in mice deficient in the RNA-editing enzyme ADAR2. Nature 406, 78–81 (2000).

4. Y. Duan, S. Dou, S. Luo, H. Zhang, J. Lu, Adaptation of A-to-I RNA editing in Drosophila. PLoS genetics 13, e1006648 (2017).

5. N. Liscovitch-Brauer et al., Trade-off between Transcriptome Plasticity and Genome Evolution in Cephalopods. Cell 169, 191–202. e111 (2017).

6. H. T. Porath et al., A-to-I RNA editing in the earliest-diverging eumetazoan phyla. Molecular biology and evolution 34, 1890–1901 (2017).

7. B. J. Liddicoat et al., RNA editing by ADAR1 prevents MDA5 sensing of endogenous dsRNA as nonself. Science 349, 1115–1120 (2015).

8. K. Pestal, C. C. Funk, J. M. Snyder, N. D. Price, P. M. Treuting, D. B. Stetson, Isoforms of RNA-editing enzyme ADAR1 independently control nucleic acid sensor MDA5-driven autoimmunity and multi-organ development. Immunity 43, 933–944 (2015).

9. N. M. Mannion et al., The RNA-editing enzyme ADAR1 controls innate immune responses to RNA. Cell reports 9, 1482–1494 (2014).

10. R. R. Kurup, E. K. Oakes, A. C. Manning, P. Mukherjee, P. Vadlamani, H. A. Hundley, RNA binding by ADAR3 inhibits adenosine-to-inosine editing and promotes expression of immune response protein MAVS. Journal of Biological Chemistry, 102267 (2022).

11. K. Niescierowicz et al., Adar-mediated A-to-I editing is required for embryonic patterning and innate immune response regulation in zebrafish. Nature communications 13, 1–14 (2022).

12. Q. Li et al., RNA editing underlies genetic risk of common inflammatory diseases. Nature, 1–9 (2022).

13. Y. Wei et al., A novel mechanism for A-to-I RNA-edited AZIN1 in promoting tumor angiogenesis in colorectal cancer. Cell death & disease 13, 1–12 (2022).

14. D. P. Reich, K. M. Tyc, B. L. Bass, C. elegans ADARs antagonize silencing of cellular dsRNAs by the antiviral RNAi pathway. Genes & development 32, 271–282 (2018).

15. P. Deng et al., Adar RNA editing-dependent and-independent effects are required for brain and innate immune functions in Drosophila. Nature communications 11, 1–13 (2020).

16. B. L. Bass, H. Weintraub, A developmentally regulated activity that unwinds RNA duplexes. Cell 48, 607–613 (1987).

17. D. Bar Yaacov, Functional analysis of ADARs in planarians supports a bilaterian ancestral role in suppressing double-stranded RNA-response. PLoS Pathog 18, e1010250 (2022).

18. D. Bar-Yaacov et al., RNA editing in bacteria recodes multiple proteins and regulates an evolutionarily conserved toxin-antitoxin system. Genome Res 27, 1696–1703 (2017).

19. J. Wolf, A. P. Gerber, W. Keller, tadA, an essential tRNA-specific adenosine deaminase from Escherichia coli. The EMBO journal 21, 3841–3851 (2002).

20. Thomas F. Wulff et al., Dynamics of diversified A-to-I editing in Streptococcus pyogenes is governed by changes in mRNA stability. Nucleic Acids Research 10.1093/nar/gkae629 (2024).

21. X. Z. Yang et al., A-to-I RNA Editing in Klebsiella pneumoniae Regulates Quorum Sensing and Affects Cell Growth and Virulence. Advanced Science, 2206056 (2023).

22. L. Didi et al., A-to-I mRNA editing in bacteria can affect protein sequence, disulfide bond formation, and function. Nucleic Acids Res 53 (2025).

23. E. Elias et al., A-to-I mRNA editing recodes hundreds of genes in dozens of species and produces endogenous protein isoforms in bacteria. Nucleic Acids Res 53 (2025).

24. D. Arad, O. Fargeon, L. Levin, S. L. Svenningsen, L. Aspit, D. Bar-Yaacov, Landscape and dynamics of TadA-dependent RNA editing in Escherichia coli reveal a role in nutrient-rich growth. mBio 10.1128/mbio.00551-26, e0055126 (2026).

25. E. Elias et al., The landscape and regulatory determinants of A-to-I RNA editing in Escherichia coli and Pseudomonas aeruginosa isolated from patients with urinary tract and ear infections. J Infect Dis 10.1093/infdis/jiaf645 (2025).

26. F. Thompson et al., Phylogeny and molecular identification of vibrios on the basis of multilocus sequence analysis. Applied and environmental microbiology 71, 5107–5115 (2005).

27. C. Baker-Austin et al., Vibrio spp. infections. Nature Reviews Disease Primers 4, 1–19 (2018).

28. F. L. Thompson, T. Iida, J. Swings, Biodiversity of vibrios. Microbiology and molecular biology reviews 68, 403–431 (2004).

29. R. Ghandour et al., ProQ-associated small RNAs control motility in Vibrio cholerae. Nucleic Acids Res 53 (2025).

30. M. Huber et al., An RNA sponge controls quorum sensing dynamics and biofilm formation in Vibrio cholerae. Nat Commun 13, 7585 (2022).

31. L. Hardy et al., The tRNA epitranscriptomic landscape and RNA modification enzymes in Vibrio cholerae. PLoS Genet 21, e1011937 (2025).

32. A. Newton, M. Kendall, D. J. Vugia, O. L. Henao, B. E. Mahon, Increasing rates of vibriosis in the United States, 1996-2010: review of surveillance data from 2 systems. Clin Infect Dis 54 Suppl 5, S391–395 (2012).

33. W. A. Norfolk, C. Shue, W. M. Henderson, D. A. Glinski, E. K. Lipp, Vibrio alginolyticus growth kinetics and the metabolic effects of iron. Microbiol Spectr 11, e0268023 (2023).

34. X. Gao et al., VqsA, a Novel LysR-Type Transcriptional Regulator, Coordinates Quorum Sensing (QS) and Is Controlled by QS To Regulate Virulence in the Pathogen Vibrio alginolyticus. Appl Environ Microbiol 84 (2018).

35. R. Lorenz et al., ViennaRNA Package 2.0. Algorithms for molecular biology 6, 1–14 (2011).

36. M. B. Miller, K. Skorupski, D. H. Lenz, R. K. Taylor, B. L. Bassler, Parallel quorum sensing systems converge to regulate virulence in Vibrio cholerae. Cell 110, 303–314 (2002).

37. W. L. Ng, L. J. Perez, Y. Wei, C. Kraml, M. F. Semmelhack, B. L. Bassler, Signal production and detection specificity in Vibrio CqsA/CqsS quorum-sensing systems. Molecular microbiology 79, 1407–1417 (2011).

38. R. C. Kelly et al., The Vibrio cholerae quorum-sensing autoinducer CAI-1: analysis of the biosynthetic enzyme CqsA. Nature chemical biology 5, 891–895 (2009).

39. D. A. Higgins, M. E. Pomianek, C. M. Kraml, R. K. Taylor, M. F. Semmelhack, B. L. Bassler, The major Vibrio cholerae autoinducer and its role in virulence factor production. Nature 450, 883–886 (2007).

40. Y. Shimony et al., M1CR0B1AL1Z3R 2.0: an enhanced web server for comparative analysis of bacterial genomes at scale. Nucleic Acids Research, gkaf413 (2025).

41. A. S. Ball, R. R. Chaparian, J. C. van Kessel, Quorum Sensing Gene Regulation by LuxR/HapR Master Regulators in Vibrios. J Bacteriol 199 (2017).

42. J. C. van Kessel, S. T. Rutherford, Y. Shao, A. F. Utria, B. L. Bassler, Individual and combined roles of the master regulators AphA and LuxR in control of the Vibrio harveyi quorum-sensing regulon. J Bacteriol 195, 436–443 (2013).

43. R. C. Kelly et al., The Vibrio cholerae quorum-sensing autoinducer CAI-1: analysis of the biosynthetic enzyme CqsA. Nat Chem Biol 5, 891–895 (2009).

44. D. Salomon et al., Type VI Secretion System Toxins Horizontally Shared between Marine Bacteria. PLoS Pathog 11, e1005128 (2015).

45. Y. Pinto, H. Y. Cohen, E. Y. Levanon, Mammalian conserved ADAR targets comprise only a small fragment of the human editosome. Genome biology 15, R5 (2014).

46. L. A. Tonkin, L. Saccomanno, D. P. Morse, T. Brodigan, M. Krause, B. L. Bass, RNA editing by ADARs is important for normal behavior in Caenorhabditis elegans. The EMBO journal 21, 6025–6035 (2002).

47. Y. A. Savva et al., Auto-regulatory RNA editing fine-tunes mRNA re-coding and complex behaviour in Drosophila. Nature communications 3, 790 (2012).

48. S. N. Deffit et al., The C. elegans neural editome reveals an ADAR target mRNA required for proper chemotaxis. Elife 6, e28625 (2017).

49. T. Ishikawa, P. K. Rompikuntal, B. Lindmark, D. L. Milton, S. N. Wai, Quorum sensing regulation of the two hcp alleles in Vibrio cholerae O1 strains. PLoS One 4, e6734 (2009).

50. K. Wu et al., CqsA-introduced quorum sensing inhibits type VI secretion system 2 through an OpaR-dependent pathway in Vibrio parahaemolyticus. Microb Pathog 162, 105334 (2022).

51. K. Kanarek, K. Keppel, H. Cohen, C. M. Fridman, M. Gerlic, D. Salomon, Assessing toxicity and competitive fitness of <i>Vibrio</i> isolates from coastal waters in Israel. mSphere 10, e00025–00025 (2025).

52. G. E. Crooks, G. Hon, J.-M. Chandonia, S. E. Brenner, WebLogo: a sequence logo generator. Genome research 14, 1188–1190 (2004).

53. M. Johnson, I. Zaretskaya, Y. Raytselis, Y. Merezhuk, S. McGinnis, T. L. Madden, NCBI BLAST: a better web interface. Nucleic acids research 36, W5–W9 (2008).

54. K. Katoh, J. Rozewicki, K. D. Yamada, MAFFT online service: multiple sequence alignment, interactive sequence choice and visualization. Briefings in bioinformatics 20, 1160–1166 (2019).

55. S. Kumar, G. Stecher, M. Li, C. Knyaz, K. Tamura, MEGA X: molecular evolutionary genetics analysis across computing platforms. Molecular biology and evolution 35, 1547–1549 (2018).

56. R. O’Toole, D. L. Milton, H. Wolf-Watz, Chemotactic motility is required for invasion of the host by the fish pathogen Vibrio anguillarum. Molecular microbiology 19, 625–637 (1996).

57. D. Salomon et al., Type VI secretion system toxins horizontally shared between marine bacteria. PLoS pathogens 11, e1005128 (2015).

58. J. Abramson et al., Accurate structure prediction of biomolecular interactions with AlphaFold 3. Nature 630, 493–500 (2024).

59. E. F. Pettersen et al., UCSF ChimeraX: Structure visualization for researchers, educators, and developers. Protein Sci 30, 70–82 (2021).

60. D. Salomon, H. Gonzalez, B. L. Updegraff, K. Orth, Vibrio parahaemolyticus type VI secretion system 1 is activated in marine conditions to target bacteria, and is differentially regulated from system 2. PLoS One 8, e61086 (2013).

61. C. M. Fridman et al., <em>Aeromonas</em> adhesins facilitate kin and non-kin attachment to enable T6SS-mediated antagonism in liquid. bioRxiv 10.64898/2026.01.27.701733, 2026.2001.2027.701733 (2026).

62. C. Camacho et al., BLAST+: architecture and applications. BMC Bioinformatics 10, 421 (2009).

